# A global baseline for qPCR-determined antimicrobial resistance gene prevalence across environments

**DOI:** 10.1101/2022.01.29.478248

**Authors:** Anna Abramova, Thomas U. Berendonk, Johan Bengtsson-Palme

## Abstract

The environment is an important component in the emergence and transmission of antimicrobial resistance (AMR). Despite that, little effort has been made to monitor AMR outside of clinical and veterinary settings. Partially, this is caused by a lack of comprehensive reference data for the vast majority of environments. To enable monitoring to detect deviations from the normal background resistance levels in the environment, it is necessary to establish a baseline of AMR in a variety of settings. In an attempt to establish this baseline level, we here performed a comprehensive literature survey, identifying 150 scientific papers containing relevant qPCR data on antimicrobial resistance genes (ARGs) in environments associated with potential routes for AMR dissemination. The collected data included 1594 samples distributed across 30 different countries and 12 sample types, in a time span from 2001 to 2020. We found that for most ARGs, the typically reported abundances in human impacted environments fell in an interval from 10^-5^ to 10^-3^ copies per 16S rRNA, roughly corresponding to one ARG copy in a thousand bacteria. Altogether these data represent a comprehensive overview of the occurrence and levels of ARGs in different environments, providing background data for risk assessment models within current and future AMR monitoring frameworks.

## Introduction

According to the World Health Organization the increasing level of antibiotic resistance over the last decades is a considerable threat to global health^1^. Already, current estimates suggest that hundreds of thousands of deaths are caused by antibiotic resistant bacteria each year worldwide, and this number is expected to reach several millions annually within the next 30 years^2^.

Antibiotic resistance has been mainly investigated in clinical and agricultural settings. However, the role of the environment as a source and dissemination route of resistance has been widely recognized in recent years^3–5^. Antibiotic resistance is common in nature and not limited to human-associated activities, as evidenced by antibiotic resistance genes (ARGs) found in glaciers, permafrost and isolated caves, which have virtually never been visited by humans^6–8^. It is even likely that resistance may be maintained independent of human activities in e.g. soil systems due to the presence of antibiotic-producing microorganisms, including certain fungal taxa^9^. Owing to the extraordinary biological diversity in nature, the external environment can serve as a source of novel ARGs not yet present in clinically relevant pathogens^10^. Furthermore, the environment constitutes a dissemination route for resistant bacteria between humans and animals^11^. However, it is not yet clear to what extent the resistant environmental bacteria and the presence of ARGs in the environment contribute to acquisition and spread of resistance in clinically relevant bacteria. To address these questions, it is essential to identify and quantify resistance in the environment. This in turn would allow pinpointing which environments possess the highest risks to human and domestic animal health and would subsequently enable us to devise measures to prevent – or at least delay – recruitment and dissemination of resistance factors from the environment.

Monitoring schemes for antibiotic resistance have been developed and put into use in clinical and agricultural settings, but systematic monitoring for antimicrobial resistance (AMR) in the environment is still lacking^12^. Existing monitoring systems largely rely on culturing of pathogens and bacterial indicator species of interest^13^, and these approaches may not be directly applicable to the monitoring of resistance in the environment. For example, the majority of environmental bacteria are not culturable using standard laboratory methods and the breakpoints for resistance are adapted for clinically relevant pathogens. Onset of molecular methods, metagenomics and quantitative real-time PCR (qPCR,) allowed to circumvent culturing drawbacks and enabled detection and quantification of individual ARGs in both culturable and unculturable bacteria. Metagenomics allows simultaneous characterization of all ARGs in an environmental sample without *a priori* knowledge, and therefore can be a promising approach for wastewater-based monitoring. However, there are several drawbacks associated with this method, including inherent variability due to lack of standardized protocols, semiquantitative nature of the generated data as well as a relatively poor quantification limit, meaning that rare genes are highly likely to be undetected. In contrast, qPCR can provide very sensitive detection and quantification of ARGs in an environmental sample, which is comparable between different studies and environmental matrices. The major hurdle for using qPCR for monitoring of AMR is the decision of which genes to select as targets in qPCR assays and what reference values to use. However, high-throughput arrays, able to quantify hundreds of ARGs in parallel have been recently developed^14^, and can partially alleviate the problem of choosing targets *a priori*^15^. Therefore, qPCR has been recognized as a promising method for AMR monitoring.

To design efficient surveillance strategies, there is a need for comprehensive background data on abundances and prevalence of ARGs occurring in both pristine and human-impacted environments. Establishing background levels of ARGs is a prerequisite for determining “safe levels” for ARGs naturally occurring in the environment and would also provide insights into to which extent an environment has been enriched with ARGs due to human activity. However, there is currently no, or very little, reference data available for the majority of environments and no agreed-upon set of genes to be used for environmental AMR monitoring.

The aim of this study was to identify a set of genes and reference abundance values in key environments, which could be used for monitoring purposes. Screening of more than 800 scientific papers for qPCR data for ARGs in various environments, we found that most surveyed environments contained a diverse set of ARGs, but generally at low abundances. It was clear that the existing data on ARG abundances is unevenly distributed across the world and that there is a general lack of data for most environments and geographical areas. Therefore, the available data could not document any significant spatiotemporal changes in AMR, but our analysis of the data identified relative abundances of ARGs that can be suggested as baseline for ARG abundances in the environment.

## Materials and Methods

### Data collection

Literature collection was performed by mining the PubMed database in April-May 2020, November 2020 and October 2021 (Figure S1). The searches were performed using several strings:

1. ‘“antibiotic resistance genes" AND (monitoring OR surveillance) AND qPCR AND’
2. ‘“antibiotic resistance genes" AND "high throughput" AND’
3. ‘"antibiotic resistance" AND (qPCR OR "quantitative PCR") AND’
4. ‘"antibiotic resistance" AND q-PCR AND‘
5. ‘"antibiotic resistance" AND "quantitative polymerase chain reaction" AND‘
6. ‘"antibiotic resistance" AND "quantitative real-time PCR" AND‘

complemented by environmental type: “sewage OR wastewater”, "surface water", sediment, soil, food, “drinking water”, “bathing water”, “swimming pools”, “bathing water”, water, airport, ("public transport" OR bus OR train OR airplane OR boat OR tram OR underground OR metro), (industrial OR pollution), abandoned, cleanroom and animal. Environmental types were selected based on the assessment of potential transmission routes for antibiotic resistance^4, 11^. This search resulted in 2636 records, and after removing duplicates in 802 unique papers. Paper suitability was assessed in several steps:

1. Initial assessment by screening the title, abstract, and methods section: review papers, methods papers, as well as papers using only metagenomics as a quantification method were discarded. Only papers using qPCR as a quantification method for environmental samples were retained. In case, papers reported experiments in artificial settings (e.g. bioreactors) as well as with addition of chemical compounds only samples that were not manipulated (e.g. control) were included in the current study.
2. In the second step, we looked through the reported results and supplementary material and selected only studies containing extractable data. Initial search revealed that collected studies used different ways of reporting ARG abundances (e.g. per 16s rRNA, per volume/weight of the sample, per total DNA or per sampled surface). We decided to retain only studies reporting gene abundances per 16S rRNA, or studies reporting numbers which could easily be converted to per 16S rRNA, to enable comparison between different environments (Figure S1). Furthermore, studies reporting only abundances per ARG class but not individual genes were also discarded.
3. For each paper the following information was extracted: first author, DOI, year, country, sample description (e.g. distance from a contamination source, season, type of fertilizer applied, stage of wastewater treatment process, source etc.) and abundances of antibiotic resistance genes (ARGs). For several studies missing information about the exact year of data collection, we extrapolated the sampling year based on the average time required from the data collection to the publication inferred from the studies reporting specific sampling dates, corresponding to an estimate of two years.
4. We divided the collected samples into several “sample types”: “air”, “biofilm”, “feces” (containing both animal and human feces), “manure”, “food”, “sediments”, “soil”, “water”, including water from different sources such as potable, surface or reclaimed water etc.; “wastewater”, consisting of wastewater, sewage and influent samples, because these samples have a similar/overlapping origin; “sludge” which includes semi-solid by-products of different stages of the wastewater treatment process; and “swab” corresponding to samples collected by swabbing the washing machines, shower drains and dishwashers surfaces with a sterile cotton swab.
5. We further classified collected samples according to exposure to anthropogenic impact: “impacted” when a sample was obviously affected by human activity such as effluent samples or wastewater, “likely impacted” when it was not directly affected by an obvious pollution source but was located in a populated area, “likely unimpacted” if the sample was collected from relatively “pristine” environments, “feces/manure” and “unknown”. The “unknown” category included samples which were clearly impacted by human activity, but where the activity aims at removing bacteria, e.g. drinking/tap water.
6. The ARG abundances were collected from text, tables and figures. Information from figures was extracted using WebPlotDigitizer v4.3 (https://automeris.io/WebPlotDigitizer). Abundances were reported as log10 ARG abundance per 16S rRNA (Supporting Information S1).

### Data analysis

Data were analyzed in RStudio v1.3.1073^16^ with the R package ggplot2 v3.3.5^17^ used to plot the graphs. To determine the abundance distributions for ARGs across environments, density plots of the ARG abundances were used, and the R summary function was used to find the quartiles for each ARG. To investigate if there were any changes in ARG levels over time, we built a linear mixed model (LMM). LMMs allow incorporation of not only fixed effects but also random effects to account for cluster-correlated data from distinct sources of variability^18^. The model was designed with ARG *Abundance* as a response variable and sampling *Year* as explanatory variable and included sampling *Country* as a random effect:

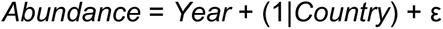

Analyses were performed using lme4 v1.1-27.1^19^ and lmerTest v3.1-3^20^ packages.

To identify a set of suitable qPCR targets, we investigated how well the data from a subset of genes could predict the abundance of the other genes in the data matrix by performing a correlation analysis. Only genes with more than 10 entries were included in this analysis. The R package stats (version 3.6.2) *cor* function with “pairwise.complete.obs” was used to calculate Spearman correlations.

## Results and Discussion

### Current data on AMR in the environment is biased

Mining of the PubMed database resulted in 802 unique papers and revealed a bias in the existing literature towards a few already fairly well studied environments (Figure S2). Unsurprisingly, the majority of the identified papers matched environmental categories that have been already extensively studied as primary sources of antibiotics and/or antibiotic resistance to the environment. These include environments linked to wastewater treatment plants, hospitals and industrial facilities, as well as agriculture and livestock production. This also highlights the undersampled nature of the other environments. Among them are categories associated with human mobility. Human mobility, in particular international travel, could significantly contribute to dissemination of ARGs across the globe^10, 21–23^. Despite that, categories associated with travel activity, such as “public transport” and “airports”, were represented by just a handful of papers. Another poorly investigated category is water associated with recreational activities. It has been suggested that accidental ingestion of water during, for example, swimming can serve as a potential route for resistant environmental bacteria into the human gut^24^.

In total, we identified 150 studies containing 1594 samples providing relevant information on levels of ARGs measured by qPCR. These samples represented 12 different sample types with “water”, “feces” and “sediments” being the most common (Figure 1B). The samples originated from 30 different countries (Figure 1A), with China being the most studied among them, both in terms of number of studies and sample types. The distribution of studies highlighted white spots on the map; in particular in South America and just a handful of studies from Africa. This is worrisome, as South America and Africa are among the regions with the highest rates of self-medication and supply of non-prescribed antibiotics in the world^25, 26^. In 2014, a WHO report on clinical data outlined Africa’s lack of established AMR surveillance systems^1^ and a review from 2017 reported that more than a third of the countries on the continent did not have recent AMR data published in the public domain, and that only little of the reported data were from surveillance^27^. The scarcity of public data on AMR in general is corroborated by the current literature survey. The majority of samples in the selected studies were collected between 2015-2018, with a gradual increase in the number of publications over the last years (Figure 1C).

**Figure 1.**
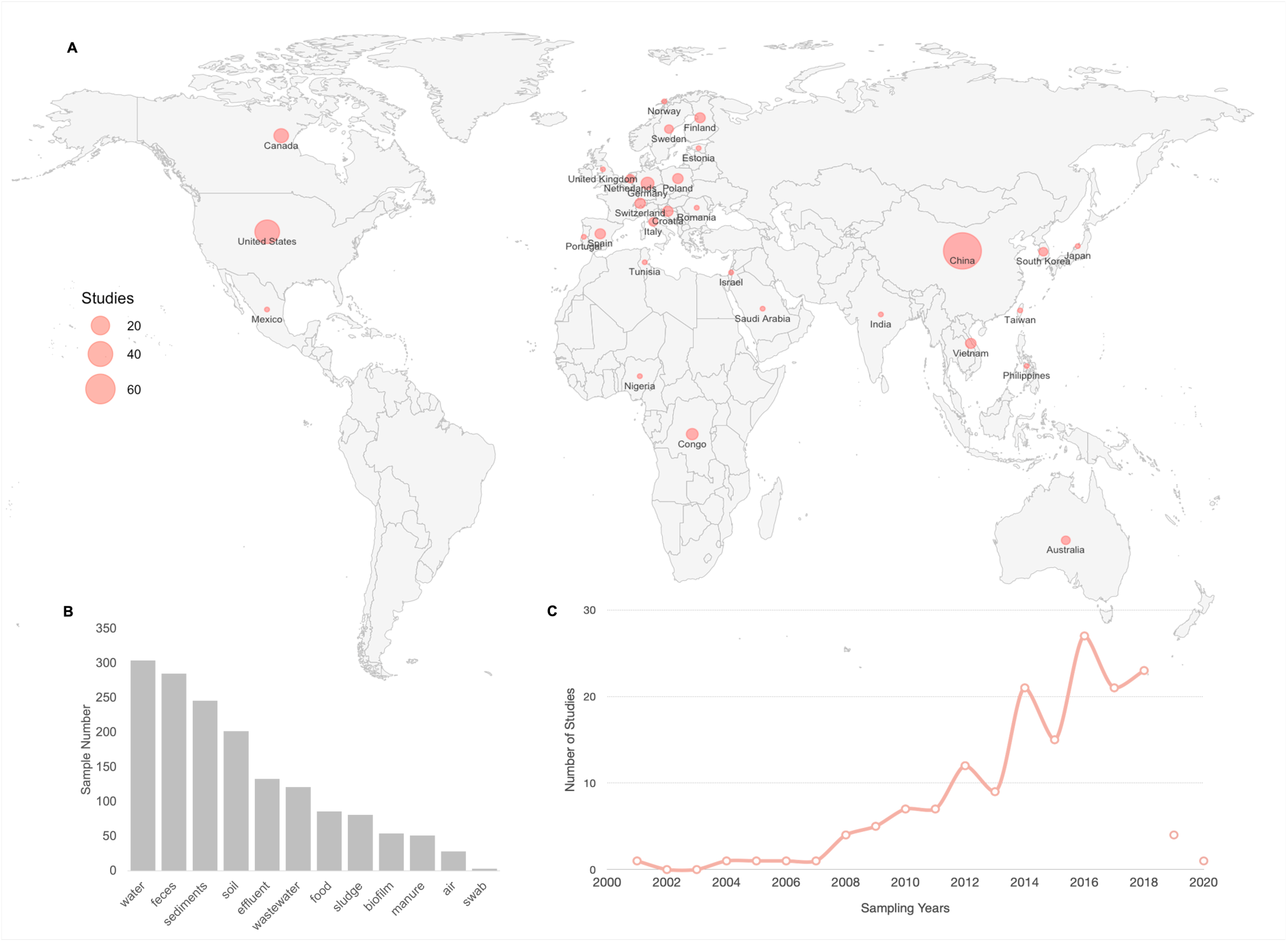
Overview of the collected data: A) geographical distribution of surveyed studies; B) number of samples representing different sample types; C) time distribution of collected studies (years 2019 and 2020 are represented by dots not connected by the line due to studies containing data from these years are yet to be published).

### Monitoring targets for AMR surveillance in the environment

It is generally agreed that the choice of targets for environmental AMR surveillance should be governed by the purpose of monitoring, which can include *i)* assessment of risks for transmission of already resistant pathogens to humans via environmental routes, *ii)* analysis of general levels of resistance in the human population, and *iii)* monitoring the evolution of resistance in the environment to identify emerging resistance threats^28^. There are several previous studies suggesting a number of ARGs targets for environmental monitoring which typically include clinically relevant ARGs to capture genes of immediate concern to human health, anthropogenic pollution markers, as well as mobility markers. The list of most often reported targets in the current survey was not very different from the previously suggested targets in Berendonk et al., 2015^4^, Bengtsson-Palme 2018^29^ and Keenum et al., 2021^30^ (Figure 2).

**Figure 2.**
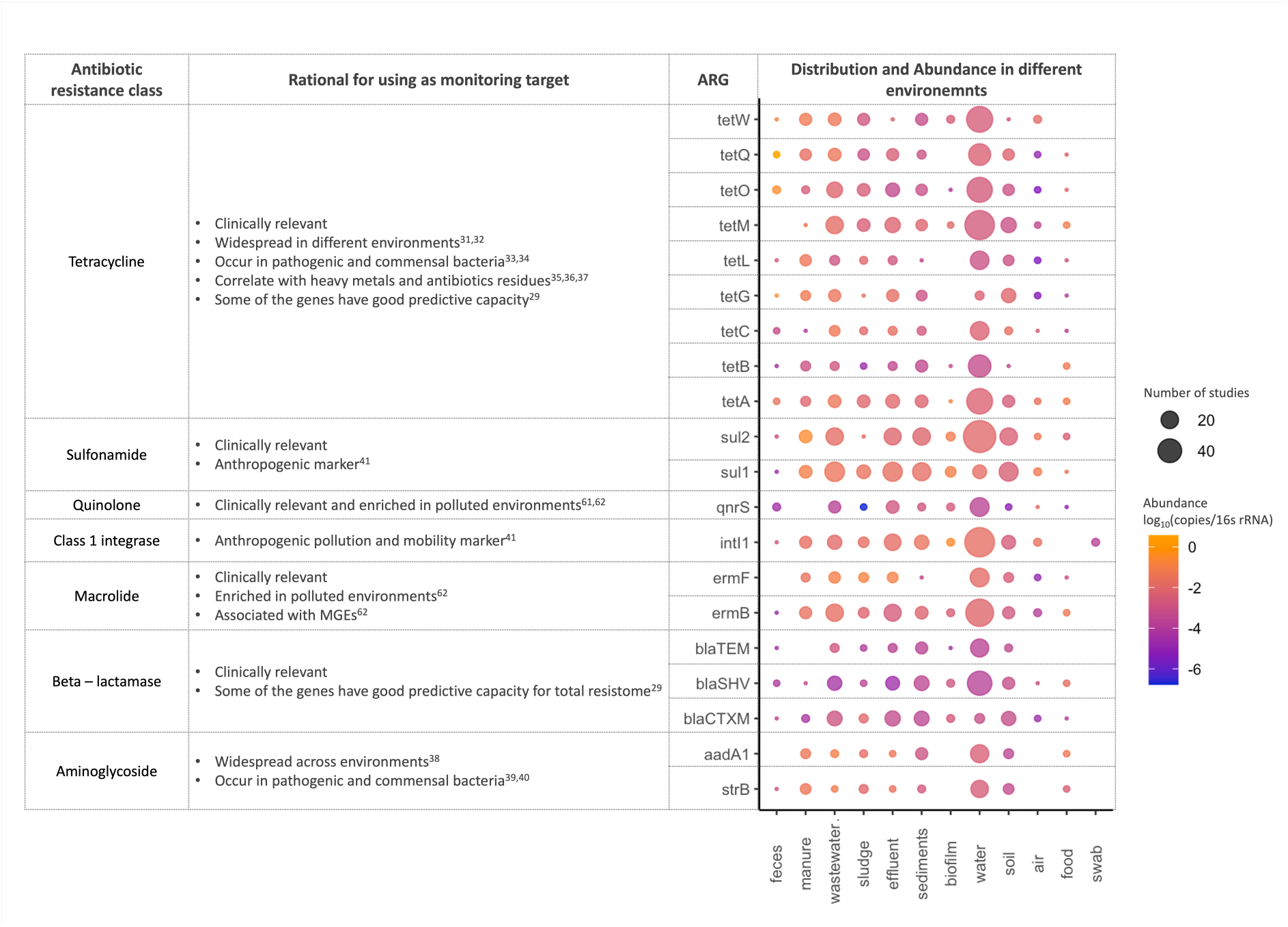
Most reported qPCR targets and their proposed value as monitoring targets suggested in existing literature; MGEs stands for Mobile Genetic Elements.

Among the ARG targets commonly used in environmental qPCR studies, many genes have already been included in clinical AMR testing (Figure 2) (e.g. https://www.oxfordbiosystems.com/AMR, https://www.opgen.com/acuitas/amr-gene-panel/) or provide resistance to antibiotics of high clinical concern listed by WHO^41^. These include clinically relevant β-lactamases such as *bla_CTX-M_*, *bla^TEM^* and *bla_SHV_*, the sulfonamide resistance gene *sul1,* as well as *ermB* and *tetM* conferring resistance to macrolides and tetracyclines, respectively. Although these genes are already common both in the environment and among human pathogens, they could be useful to predict general resistance levels or transmission risks. However, their usefulness in environmental monitoring for emerging threats is somewhat questionable. At the same time, they may function as a gauge of the total antibiotic resistance content in an environment, as some of them have been shown to be good predictors of the total resistome, particularly *tetM* and *bla_TEM_*^29^.

Not typically included in routine clinical monitoring but previously suggested as a marker for anthropogenic AMR dissemination^42^ in the environment is the class 1 integrase *intI1* gene. Despite that this gene is not a resistance gene itself, it is often linked to ARGs. According to the existing literature, it is widespread in the environment, found in both pathogenic and commensal bacteria, enriched by anthropogenic activities and correlates well with the abundances of other commonly occurring ARGs^43^. Interestingly, despite being so widespread, our data showed that this gene has lower abundances in the likely unimpacted sampling locations, such as arctic soils and alpine lakes, supporting its usefulness as an indicator of anthropogenic pollution in the environment. Similarly, the sulfonamide resistance gene *sul2* has also been suggested as a monitoring target by Berendonk et al. (2015)^4^, was among the most reported in our study and was widespread across all the environments. However, *sul2* added little information compared to *intI1* (Figure S3), so these two genes would be mostly redundant in AMR monitoring.

The plasmid-mediated fluoroquinolone resistance gene *qnrS* was among the most reported genes in our study. It was found in all environments except manure and swabs. Contrary to what has been reported previously^61, 62^, we did not detect an enrichment of this gene in polluted environment specifically. As it is the only commonly reported fluoroquinolone ARG, it is an interesting target for monitoring. However, it has the drawback of generally not being able to induce clinically relevant levels of resistance without additional resistance mechanisms, which makes its clinical impact somewhat limited.

Among the most common target genes, several tetracycline resistance genes showed significantly higher abundances in fecal/manure samples than in other human-impacted environments (including *tetA*, *tetG*, t*etH*, *tetO*, *tetQ* and *tetW*). Furthermore, these genes are among the most powerful ARGs for predicting the diversity and abundance of other ARGs^29^ and can therefore, despite their widespread distribution, still be useful in monitoring to extrapolate other parts of the resistome.

There were also several genes that were not often included in qPCR studies, but when they were, they often appeared in relatively high abundances (above one copy per 1000 bacteria) (Figure S5). Such ARGs could be interesting additional monitoring targets in future AMR monitoring schemes. Among these genes was the vancomycin resistance gene *vanA,* a previously suggested indicator of antibiotic resistance contamination of clinical origin, which is thought to be uncommon in the environment^44^. In our data, however, *vanA* was about as abundant as other ARGs in the environment when it has been looked for. Another gene that seemed to correlate with anthropogenic activities is *ereA*, a macrolide resistance gene which has previously been reported to be the most abundant in metal polluted soil^45^ and is enriched by long-term application of manure^46^. In contrast, the *mexF* gene has been reported from many different environments, including soil, sediments and water^21^ and is suggested to naturally occur in unaffected environments such as pristine Antarctic soils^47^, indicating that it might be a useful target for identifying enrichment of ARGs occurring in environmental microbial communities irrespectively of pollution from fecal material. The *sul3* gene has been rarely reported, despite being rather abundant in the cases when it was quantified. According to previous studies, *sul3* is typically less frequent and abundant than *sul1* and *sul2*^31, 48^. Similarly to *sul1* and *sul2*, it has been shown to be associated with class 1 integrons and has been detected on a conjugative plasmid, suggesting its potential to be horizontally transferred and disseminated in the environment^49^. Interestingly, *sul3* was first detected in an *E. coli* isolate from a pig, and it was later found in both healthy and diseased humans. Furthermore, it is often enriched in polluted environments and is present in both commensal and pathogenic bacteria^50^. Several other genes (*tnpA4, tnpA5, qacEdelta, aadA2, floR, tet32, cmlA1* and *mefA*), which were not often reported, but often highly abundant when detected, would also be potentially suitable additional targets for AMR monitoring (Figure S5).

### Around one in a thousand environmental bacteria carry clinical ARGs

In the current study, we estimated typical abundance levels of ARGs in different environments. We found that for most ARGs, the typically reported abundances fell in an interval from 10^-5^ to 10^-3^ copies per 16S rRNA, roughly corresponding to one ARG copy in a thousand bacteria (Figure 3B). Importantly, there was a strong bias towards studies of environments already impacted by humans (e.g. WWTPs, agriculture and animal production), and therefore this range rather reflects ARG levels in environments already affected by human activity.

**Figure 3.**
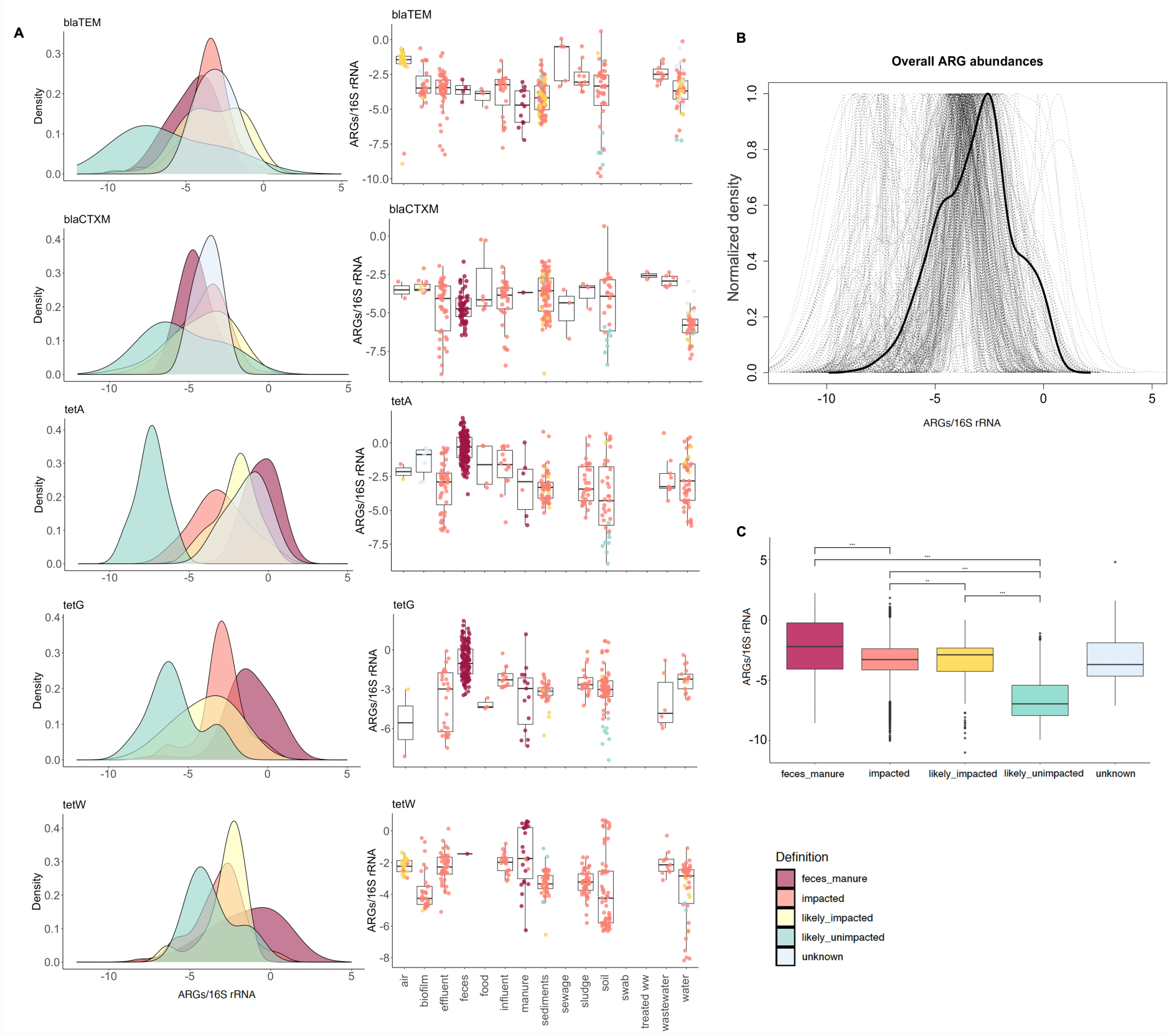
Relative abundance distributions of surveyed ARGs. A) Density plots represent abundance ranges of blaTEM, blaCTXM, tetA, tetG and tetW, in different environmental types divided according to exposure to anthropogenic impact (see material and methods for the detailed description), and corresponding bar plots show what samples contributed to each distribution. B) Density plot representing relative abundance distributions across all the studied genes with the bold line indicating the average distribution. C) Difference between abundance ranges in different environmental types. The environmental types are color-coded as follows: “feces/manure” in purple, “impacted” in orange, “likely impacted” in yellow, “likely unimpacted” in green and “unknown” type in blue. Note that abundances are given relative to the number of 16S rRNA copies, which means that these distributions do not necessarily correspond to the total exposure to resistant bacteria in a given environment.

That said, most of the ARGs measured in likely unimpacted sampling locations had lower typical abundances, below 10^-5^ copies per 16S rRNA (Figure 3A). In particular, samples of water and soil from the arctic tundra showed the lowest ARG abundances. There are previous studies suggesting that ARG abundances fluctuate with latitude, which to some degree could be explained by optimal growth temperature for the microbes carrying these ARGs^51^. Furthermore, there is evidence for a distance-decay relationship for abundance of ARGs on the global and continental scales^52, 53^, which could explain higher abundances of ARGs in the water from alpine lakes in comparison to water from arctic tundra. Overall, however, it is hard to draw any major conclusions due to the small number of samples from relatively unimpacted locations.

Our data revealed distinct distributions of several tetracycline genes (including *tetA*, *tetG*, t*etH*, *tetO*, *tetQ* and *tetW*) in feces/manure samples (Figure 3A, Figure S6). Tetracycline ARGs constituted the predominant class among the most reported qPCR targets and are typically associated with human fecal samples^54^. Notably, several of these genes showed distinct abundance distributions for impacted environments and feces/manure samples, suggesting that they can be used as markers for human/animal fecal contamination in the environment. Particularly, *tetA* and *tetG* show distinct distributions in impacted and non-impacted environments and could serve as indicators for anthropogenic pollution, while *tetW* shows overlapping distributions making it less useful to ascertain human impact. Similarly, the clinically relevant *bla_CTX-M_* and *bla_TEM_* genes also showed a separation in impacted and non-impacted environments, although that was less clear than for tetracycline genes.

The “unknown” category was characterized by relatively high abundances of ARGs and for some of the genes even higher levels than in impacted samples. This category comprised samples of water and biofilm from households (unspecified), as well as potable and reclaimed water distribution systems. We classified these samples as “unknown” since they are obviously impacted by human activity, but the impact targets the removal of ARGs and bacteria. Despite that, ARGs in some of these samples had higher relative abundances than in the fecal/manure samples (Figure 3A, Figure S6). These samples may yet not have higher abundances of ARGs per amount of sample, as this would also depend on how much bacteria are present in the samples per volume or weight.

### It is unclear whether ARG abundances have increased over time

To investigate if there were changes in ARG abundances over time, we used linear mixed models on gene abundances from each of the environments, accounting for variability between countries (Figure S4, Table S2). Due to the successively increased sensitivity of the methods used for qPCR and inclusion of a larger number of ARGs profiled in each study over time, we chose to use linear mixed models for the maximum values per year reported from a particular sample type rather than all ARG values reported. Using this approach, we found positive trends for most of the environments (for cases where there was enough data to produce the estimates), except for in effluent, sediments, sludge, wastewater and water where the trends were negative (Table S2). That said, the trends were significant only for biofilm, effluent, food, sediments, sludge and wastewater. Thus, the most prominent finding of this analysis might be that in most cases there was too little data to draw any conclusions on ARG abundance changes over time, highlighting the need for time series data to understand the long-term development of antibiotic resistance in the environment.

### Abundance data for some ARGs provide redundant information

The set of most reported genes consists of potential candidates to be used as qPCR targets, since they were on average more abundant and were also found in most environments (Figures 2 and 3). To explore if they would be good predictors of overall ARG abundances, we performed a correlation analysis of the most reported genes and the rest of the genes in the data matrix (Figure S3). This analysis revealed a certain degree of redundancy in terms of what information can be gained from the proposed monitoring targets. ARGs proposed as monitoring targets that strongly correlate with each other convey similar information on ARG abundance and diversity and may therefore not be very useful to use in the same panel for ARG monitoring in the environment. Genes that were overall redundant included the *tetO* and *tetA* genes, *strB* and *ermF*, *bla_CTX-M_* and *bla_SHV_*, *tetG* and *tetM*, as well as *intI1* and *sul2* (Figure S3). If resources are constrained, it would seem wise not to use several genes in these smaller groups together in the same ARG panel for monitoring.

### Specific uses of qPCR as a monitoring tool

In this study, we have exclusively focused on the use of qPCR as a tool for monitoring the abundance of ARGs in the environment. In general, the qPCR technique has several benefits, but also some drawbacks as a surveillance tool. An important advantage is that it is a highly sensitive method that can detect much lower levels of ARGs than is currently possible using shotgun metagenomic sequencing^55^. Furthermore, it can operate on very minute quantities of DNA, which makes it suitable also for low-biomass samples. In addition, as it is performed on DNA, it can detect ARGs in non-culturable bacteria and is functional also on complex samples with many different species.

However, application of qPCR as an AMR monitoring tool requires *a priori* knowledge on a set of targets as well as how to interpret the results in terms of when an ARG occurrence pattern becomes a concern. The limiting factor of predefined targets can be partially overcome by using qPCR arrays with hundreds of genes^15^, but this also comes with increased costs and may not (at present) be feasible for large-scale routine monitoring of environmental AMR. Despite that, even a handful of selected targets can still provide useful information about the total resistome situation, fecal contamination and potentially HGT intensity. From a monitoring perspective, the background ARG levels identified in this study could be used to infer an increase in ARG abundances as a sign of pollution and/or selection. Based on the collected data, we advise the use of 10 times the third quartile (3Q) values for a given ARG (Table S1) to determine the upper limit for what should be viewed as a deviation from normal background levels in the environment. A more fundamental aspect of such deviations is whether an increase in the abundance of ARGs or mobile genetic elements in a particular environment or at a specific time point is a relevant indicator of a selective pressure for resistance. Importantly, an increase of ARG abundances without any context is more likely to be an indicator of human pollution^56^, but it cannot be ruled out that such a change could be due to a specific selection pressure from antibiotics^57^, by co-selection from other antibacterial compounds^58^, or simply from taxonomic shifts that are unrelated to antibiotic resistance.

In the end, the suitability of qPCR for environmental AMR surveillance comes down to the purpose of monitoring and what type of actions one might want to take based on the monitoring results. If detecting dissemination of known high-risk ARGs is the sole purpose of monitoring, qPCR is an excellent method thanks to its sensitivity. However, if identification of emergent resistance threats is the goal, qPCR is unlikely to give useful guidance; instead, shotgun metagenomics^59^ or selective culturing followed with genetic profiling would provide more useful information.

### Outcomes and recommendations

In this study, we performed a literature survey to explore the abundance and prevalence of ARGs in various environments as quantified by qPCR. We found that, overall, previous suggestions for ARGs to be included in environmental AMR monitoring^4, 29^ seem relevant. Particularly, inclusion of the *intI1*, *sul1*, *bla_TEM_*, *bla_CTX-M_* and *qnrS* genes in environmental monitoring seems essential, along with a selection of the tetracycline genes, in particular at least one of *tetA* or *tetG,* which could serve as fecal pollution markers. However, there are also genes that are not often looked for in qPCR surveys that perhaps should be, including *sul3*, *vanA*, *tetH*, *aadA2*, *floR*, *ereA* and *mexF*. These genes are abundant in some environments, but were not often included in qPCR studies of environmental AMR. We also provide environmental baseline levels for the ARGs studied through qPCR (Figure 3, Table S1); for most ARGs the typical relative abundance falls in an interval from 10^-5^ to 10^-3^. It should be noted that this is the range of normal abundances of ARGs and should not be considered a maximum acceptable limit of these ARGs in any given environment. Such maximum acceptable levels need to be determined taking risks to human health (and potentially the environment) as well as the numbers of bacteria in a given volume of sample into account^5, 60^, and would also need to consider transmission routes to humans^11^. The different standards of reporting DNA abundances constituted a complicating factor for this study. Particularly, the choice to report the abundances of ARGs relative to the 16S rRNA gene, which was necessitated by the availability of data, limits the interpretability of our data in terms of exposure risks. The absence of clear trends of increases or decreases in ARG abundances over time indicates a need for more systematic time series data on ARG abundances in a variety of environments. Our results also highlight the scarcity of AMR data from parts of the world, particularly from Africa and South America, and underscores the need for a concerted effort to quantify typical background levels of AMR in the environment more broadly to enable efficient environmental AMR surveillance schemes akin to those that exist in clinical settings.

## Acknowledgments

JBP acknowledges funding from the Swedish Research Council (VR; grant 2019-00299) under the frame of JPI AMR (EMBARK; JPIAMR2019-109), the Data-Driven Life Science (DDLS) program supported by the Knut and Alice Wallenberg Foundation (KAW 2020.0239), the Swedish Foundation for Strategic Research (FFL21-0174), the Centre for Antibiotic Resistance Research at the University of Gothenburg, the Sahlgrenska Academy at the University of Gothenburg, and the Swedish Cancer and Allergy fund (Cancer-och Allergifonden). TUB acknowledges funding of the JPI AMR – EMBARK project funded by the Bundesministerium für Bildung, und Forschung (BMBF) under grant number F01KI1909A. We would like to thank Marcus Wenne, Emil Burman, Sebastian Wettersten, Tora Hulterström and anonymous reviewers for constructive feedback on earlier versions of the manuscript.

