## Supplementary Material for "A global baseline for qPCR-determined antimicrobial resistance gene prevalence across environments"

### Supporting Information

**Figure S1. Literature survey PRISMA flow diagram.**

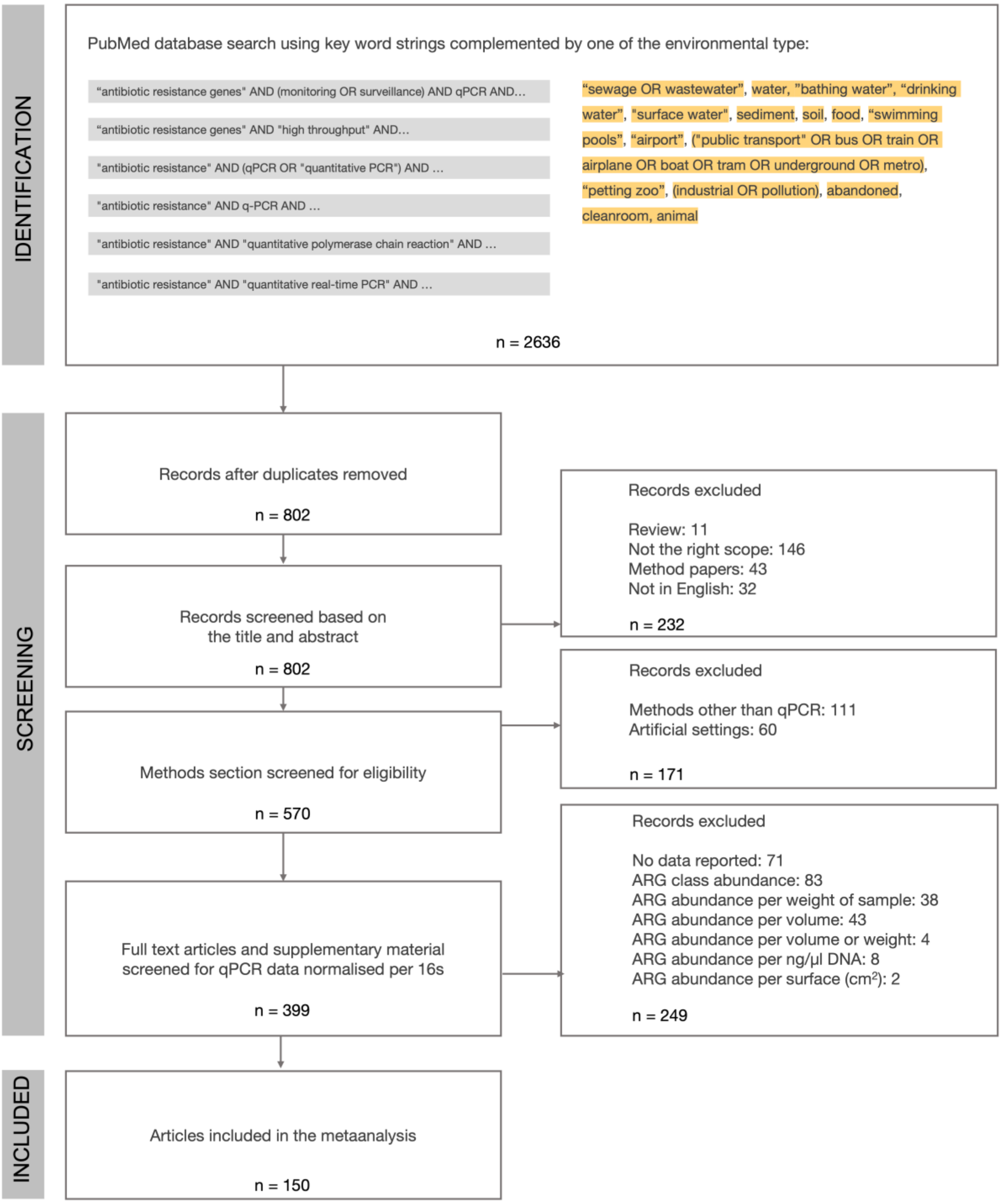

**Figure S2. Overview of the PubMed search results stratified according to different search terms.**

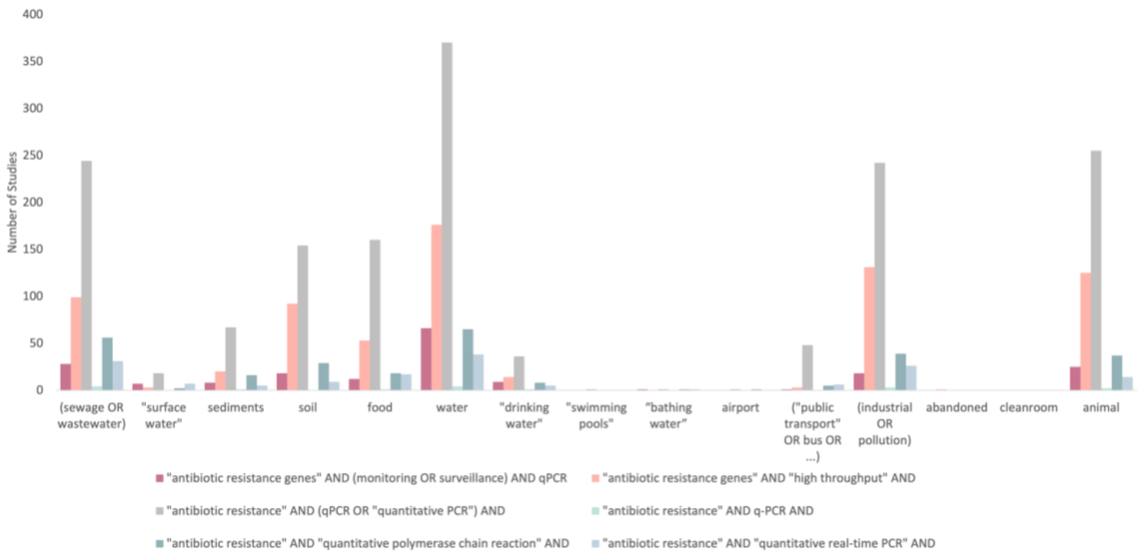


**Table S1. Abundance distribution for reported ARGs**

| Gene | Observations | Min | 1Q | Median | Mean | 3Q | Max |
|---|---|---|---|---|---|---|---|
| aac | 30 | -5.465 | -4.691 | -4.316 | -4.088 | -3.862 | -0.448 |
| aac6I | 13 | -5.115 | -4.458 | -4.313 | -4.230 | -3.786 | -3.546 |
| aac6Ib | 105 | -9.392 | -3.747 | -3.129 | -3.499 | -2.399 | 0.833 |
| aac6II | 53 | -9.348 | -4.239 | -3.527 | -3.826 | -2.991 | -1.450 |
| aac6Iy | 6 | -5.542 | -5.325 | -5.207 | -5.037 | -4.986 | -3.993 |
| aac2ic | 8 | -9.650 | -8.451 | -8.316 | -8.342 | -8.102 | -7.458 |
| aac3ia | 7 | -7.932 | -7.736 | -4.387 | -4.643 | -1.645 | -1.419 |
| aac3iv | 6 | -8.919 | -8.112 | -7.471 | -7.643 | -7.204 | -6.554 |
| aac3vi | 8 | -7.317 | -6.171 | -5.668 | -5.698 | -5.322 | -3.939 |
| aac6Ia | 6 | -8.757 | -8.438 | -8.027 | -7.945 | -7.479 | -6.987 |
| aacA | 26 | -4.398 | -4.187 | -3.704 | -3.516 | -3.157 | -1.658 |
| aacC | 81 | -6.246 | -3.552 | -2.536 | -2.983 | -1.999 | -1.424 |
| aacC1 | 8 | -4.648 | -4.484 | -3.568 | -3.787 | -3.247 | -2.998 |
| aacC2 | 33 | -5.127 | -3.993 | -3.589 | -3.455 | -2.745 | -1.437 |
| aacC4 | 43 | -4.835 | -3.856 | -3.484 | -3.270 | -2.860 | -0.452 |
| aadA1 | 146 | -8.492 | -3.471 | -2.792 | -2.924 | -2.008 | -0.150 |
| aadA2 | 110 | -5.542 | -3.665 | -2.802 | -2.910 | -2.025 | -0.893 |
| aadA5 | 73 | -8.257 | -3.894 | -3.336 | -3.679 | -2.739 | -1.455 |
| aadA9 | 57 | -8.127 | -5.385 | -4.177 | -4.510 | -3.668 | -1.958 |
| aadD | 18 | -4.914 | -4.223 | -3.613 | -3.429 | -2.518 | -1.187 |

| | | | | | | | |
|---|---|---|---|---|---|---|---|
| aadE | 49 | -4.349 | -3.745 | -3.552 | -3.410 | -2.968 | -1.693 |
| acrA1 | 66 | -9.798 | -4.774 | -3.677 | -4.112 | -3.274 | -2.390 |
| acrA2 | 37 | -5.242 | -3.846 | -3.396 | -3.357 | -2.870 | -1.242 |
| acrA3 | 18 | -5.576 | -4.632 | -4.298 | -4.328 | -3.933 | -3.358 |
| acrA4 | 57 | -4.987 | -3.562 | -3.137 | -3.205 | -2.854 | -1.315 |
| acrA5 | 55 | -4.889 | -3.564 | -3.171 | -3.191 | -2.867 | -1.599 |
| acrB | 31 | -9.378 | -5.179 | -3.373 | -4.111 | -2.475 | -0.140 |
| acrD | 2 | -7.619 | -7.554 | -7.488 | -7.488 | -7.422 | -7.357 |
| acrF | 32 | -8.684 | -5.433 | -4.175 | -4.702 | -3.277 | -2.593 |
| acrR1 | 45 | -8.788 | -4.660 | -3.757 | -4.356 | -3.447 | -2.510 |
| acrR2 | 23 | -5.554 | -4.355 | -3.394 | -3.158 | -1.966 | -0.554 |
| adeA | 17 | -8.428 | -7.613 | -4.488 | -5.336 | -3.654 | -3.136 |
| aerA3 | 2 | -4.256 | -4.054 | -3.852 | -3.852 | -3.650 | -3.448 |
| blaampC | 94 | -11.000 | -4.130 | -3.543 | -3.838 | -3.262 | -0.690 |
| ant2ia | 5 | -8.140 | -7.876 | -7.625 | -7.500 | -7.545 | -6.313 |
| ant3ia | 6 | -9.348 | -8.446 | -8.165 | -8.027 | -7.828 | -6.228 |
| aph | 5 | -4.222 | -3.894 | -3.846 | -3.016 | -1.726 | -1.389 |
| aph2Id | 5 | -4.502 | -2.663 | -2.575 | -2.846 | -2.350 | -2.139 |
| aph3ia | 1 | -1.012 | -1.012 | -1.012 | -1.012 | -1.012 | -1.012 |
| aph6Ia | 6 | -8.137 | -7.596 | -5.369 | -5.235 | -2.770 | -2.295 |
| aphA1 | 52 | -5.845 | -4.209 | -3.474 | -3.475 | -2.879 | -1.258 |
| aphA3 | 54 | -5.736 | -4.694 | -3.714 | -3.756 | -3.187 | -0.304 |
| bacA1 | 21 | -7.861 | -6.455 | -4.458 | -4.924 | -3.699 | -2.629 |
| bacA2 | 29 | -5.252 | -4.280 | -3.445 | -3.673 | -3.267 | -2.505 |
| blaAAC | 14 | -9.253 | -7.685 | -4.423 | -5.294 | -4.276 | -0.974 |
| blaCMY2 | 57 | -8.566 | -4.213 | -3.678 | -3.891 | -3.267 | -0.280 |
| blaEC | 7 | -9.869 | -8.522 | -7.441 | -7.929 | -7.405 | -6.337 |
| blaNDM | 58 | -7.861 | -6.336 | -5.680 | -5.505 | -4.764 | -2.277 |
| blaMOX | 19 | -8.813 | -4.341 | -3.685 | -3.879 | -2.952 | -2.537 |
| blaOCH | 25 | -7.944 | -4.699 | -4.228 | -4.627 | -3.824 | -3.323 |
| blaPAO | 8 | -9.334 | -7.836 | -4.883 | -5.694 | -4.130 | -2.543 |
| blaSM | 5 | -6.979 | -6.896 | -6.861 | -6.751 | -6.764 | -6.255 |
| blaROB | 22 | -8.521 | -7.308 | -4.613 | -5.538 | -4.308 | -3.182 |
| blaTEM | 376 | -9.826 | -4.405 | -3.485 | -3.665 | -2.678 | 0.600 |
| blaCTXM | 335 | -8.992 | -5.406 | -4.040 | -4.312 | -3.199 | 0.642 |
| blaOXY1 | 64 | -7.924 | -3.821 | -3.226 | -3.412 | -2.639 | -2.126 |
| blaSHV2 | 16 | -8.346 | -6.853 | -4.171 | -4.971 | -3.432 | -3.130 |
| blaPSE1 | 24 | -9.749 | -4.622 | -3.906 | -4.509 | -3.368 | -1.742 |
| blaAXA1 | 2 | -8.759 | -8.627 | -8.496 | -8.496 | -8.364 | -8.233 |
| blacepA | 24 | -7.668 | -5.375 | -5.266 | -5.029 | -4.836 | -3.187 |
| blacfxA | 18 | -8.242 | -6.721 | -4.285 | -4.633 | -2.967 | -1.371 |
| blacphA | 33 | -8.394 | -5.590 | -4.742 | -4.782 | -3.565 | -2.140 |
| blaIMP | 44 | -8.772 | -3.934 | -3.511 | -3.326 | -2.319 | -0.319 |

| | | | | | | | |
|---|---|---|---|---|---|---|---|
| blaVIM | 33 | -9.914 | -4.912 | -4.156 | -4.223 | -2.250 | -1.500 |
| blaOXA2 | 14 | -3.854 | -3.247 | -2.648 | -2.743 | -2.393 | -1.460 |
| blaOXA48 | 36 | -8.142 | -5.791 | -4.691 | -4.479 | -2.490 | -1.706 |
| blaOXA58 | 62 | -8.000 | -5.387 | -3.031 | -3.805 | -2.394 | -0.897 |
| blaL1 | 26 | -4.969 | -4.190 | -3.841 | -3.822 | -3.520 | -2.463 |
| bla1 | 7 | -4.339 | -3.763 | -3.527 | -3.552 | -3.176 | -3.123 |
| blaGES | 50 | -8.135 | -4.379 | -3.261 | -3.551 | -2.430 | -1.198 |
| blaCMY | 18 | -4.743 | -4.224 | -3.907 | -3.844 | -3.441 | -2.781 |
| blaI | 9 | -8.800 | -8.322 | -7.629 | -7.530 | -7.199 | -4.333 |
| blaKPC | 47 | -6.796 | -4.483 | -3.993 | -3.858 | -2.920 | -1.464 |
| blaM | 24 | -4.242 | -3.473 | -3.207 | -2.765 | -2.613 | 0.249 |
| blaOKP | 6 | -5.242 | -4.091 | -3.815 | -4.002 | -3.716 | -3.273 |
| blaOXA1 | 95 | -8.085 | -3.981 | -3.121 | -3.354 | -2.509 | -0.855 |
| blaPER | 24 | -4.977 | -4.335 | -4.174 | -3.974 | -3.596 | -3.123 |
| blaS1 | 10 | -3.587 | -3.358 | -3.165 | -3.143 | -2.885 | -2.745 |
| blaSFO | 87 | -6.354 | -4.704 | -3.297 | -3.778 | -2.828 | -1.959 |
| blaSHV | 140 | -7.946 | -5.063 | -3.651 | -3.902 | -2.750 | 0.059 |
| blaSHV34 | 8 | -2.654 | -2.560 | -2.430 | -2.170 | -2.137 | -0.542 |
| blaTLA | 23 | -5.200 | -3.729 | -3.040 | -3.338 | -2.900 | -2.260 |
| blaVEB | 25 | -4.398 | -3.724 | -3.237 | -2.915 | -2.079 | -0.975 |
| blaZ | 11 | -5.582 | -5.291 | -4.876 | -5.042 | -4.744 | -4.674 |
| cadA | 15 | -7.619 | -5.165 | -4.860 | -5.176 | -4.615 | -4.140 |
| carBR | 7 | -9.129 | -8.895 | -8.547 | -8.430 | -8.343 | -6.858 |
| catA1 | 25 | -4.596 | -4.127 | -3.776 | -3.315 | -2.682 | -1.373 |
| catB3 | 32 | -9.572 | -4.050 | -3.252 | -3.893 | -2.420 | -0.980 |
| catB8 | 25 | -8.712 | -3.800 | -3.417 | -3.821 | -3.049 | -2.464 |
| qacEdelta | 131 | -8.845 | -3.439 | -2.314 | -2.711 | -1.716 | -0.014 |
| ceoA | 61 | -8.240 | -3.956 | -3.618 | -3.965 | -3.251 | -2.773 |
| cepA | 9 | -4.567 | -4.000 | -3.753 | -3.806 | -3.517 | -3.390 |
| cfiA | 12 | -8.944 | -5.752 | -4.559 | -5.365 | -4.006 | -3.699 |
| cfr | 19 | -5.045 | -1.600 | -0.076 | -1.116 | 0.059 | 0.385 |
| cfxA | 40 | -5.515 | -3.858 | -3.573 | -3.454 | -3.169 | -0.824 |
| chrA | 2 | -7.176 | -6.995 | -6.814 | -6.814 | -6.633 | -6.452 |
| cIntI1 | 7 | -1.771 | -1.402 | -1.236 | -1.195 | -0.987 | -0.580 |
| cmeA | 2 | -5.479 | -5.027 | -4.575 | -4.575 | -4.123 | -3.671 |
| cmeAF | 1 | -3.863 | -3.863 | -3.863 | -3.863 | -3.863 | -3.863 |
| cmle3 | 7 | -9.496 | -7.854 | -6.635 | -7.251 | -6.560 | -5.796 |
| cmlA1 | 84 | -8.474 | -3.849 | -2.950 | -2.905 | -1.891 | 0.385 |
| cmr | 16 | -6.732 | -6.034 | -5.234 | -4.903 | -4.330 | -1.738 |
| cmxA | 54 | -6.452 | -4.328 | -3.773 | -3.765 | -3.344 | -1.395 |
| copA | 1 | -0.255 | -0.255 | -0.255 | -0.255 | -0.255 | -0.255 |
| cphA | 69 | -4.929 | -3.629 | -3.270 | -3.265 | -2.814 | -1.182 |
| crcA | 1 | -0.141 | -0.141 | -0.141 | -0.141 | -0.141 | -0.141 |

| | | | | | | | |
|---|---|---|---|---|---|---|---|
| dfrA1 | 73 | -9.349 | -4.198 | -3.257 | -3.574 | -2.463 | -0.906 |
| dfrA12 | 19 | -4.728 | -4.245 | -3.985 | -3.934 | -3.718 | -2.899 |
| dfrA14 | 8 | -3.000 | -2.395 | -2.010 | -1.968 | -1.600 | -0.980 |
| dfrA22 | 9 | -5.155 | -5.000 | -3.853 | -4.076 | -3.283 | -3.174 |
| emrD | 58 | -9.274 | -4.886 | -3.334 | -3.824 | -2.574 | -1.729 |
| ereA | 92 | -5.754 | -3.986 | -3.173 | -2.950 | -1.959 | 1.000 |
| ereB | 30 | -8.848 | -4.054 | -3.660 | -4.194 | -3.296 | -2.443 |
| erm34 | 13 | -8.799 | -7.670 | -4.537 | -5.446 | -4.177 | -2.890 |
| erm35 | 14 | -5.543 | -4.637 | -3.578 | -3.893 | -3.288 | -2.527 |
| erm36 | 57 | -8.221 | -5.199 | -3.839 | -4.369 | -3.294 | -1.566 |
| ermA | 35 | -8.800 | -4.410 | -3.342 | -3.668 | -2.584 | -1.390 |
| ermB | 332 | -8.992 | -3.431 | -2.554 | -2.744 | -1.769 | 0.962 |
| ermC | 73 | -7.985 | -4.771 | -3.141 | -3.755 | -2.495 | -0.080 |
| ermF | 125 | -7.770 | -2.883 | -2.311 | -2.345 | -1.269 | -0.046 |
| ermJ.ermD | 13 | -3.969 | -3.472 | -3.262 | -3.314 | -3.164 | -2.726 |
| ermK | 11 | -9.887 | -6.035 | -4.468 | -5.275 | -4.054 | -2.994 |
| ermT | 32 | -4.764 | -4.197 | -3.925 | -3.675 | -3.548 | -0.710 |
| ermX | 67 | -7.268 | -4.424 | -3.359 | -3.539 | -2.387 | -1.580 |
| ermY | 9 | -4.347 | -4.117 | -3.939 | -3.893 | -3.817 | -2.947 |
| fabK | 5 | -9.264 | -8.513 | -8.388 | -7.038 | -4.626 | -4.398 |
| fexB | 12 | -3.825 | -0.352 | -0.246 | -0.465 | -0.118 | 0.625 |
| floR | 113 | -6.194 | -4.447 | -3.394 | -3.141 | -2.261 | 0.621 |
| folA | 10 | -5.610 | -4.933 | -4.272 | -4.389 | -3.798 | -3.360 |
| fox5 | 83 | -9.375 | -4.909 | -3.246 | -3.854 | -2.722 | -1.719 |
| gadAB | 5 | -4.716 | -4.467 | -3.755 | -4.038 | -3.658 | -3.594 |
| ImrA | 4 | -4.323 | -4.247 | -3.945 | -3.952 | -3.649 | -3.593 |
| IncWrepA | 17 | -6.521 | -4.814 | -4.435 | -4.452 | -3.502 | -2.794 |
| intI1 | 543 | -8.992 | -3.136 | -2.259 | -2.366 | -1.343 | 1.600 |
| intI2 | 25 | -6.883 | -4.913 | -4.172 | -3.877 | -2.845 | -0.478 |
| intI3 | 17 | -6.926 | -5.145 | -3.459 | -3.626 | -2.634 | -0.690 |
| lnuB | 37 | -8.398 | -3.578 | -3.254 | -3.361 | -2.795 | -1.944 |
| IS613 | 48 | -6.442 | -4.469 | -3.594 | -3.485 | -2.469 | -0.714 |
| korB | 21 | -4.240 | -3.013 | -2.495 | -2.666 | -2.190 | -1.467 |
| lmrA | 1 | -8.231 | -8.231 | -8.231 | -8.231 | -8.231 | -8.231 |
| lnuA | 27 | -6.000 | -4.419 | -3.854 | -3.963 | -3.519 | -2.278 |
| lnuC | 24 | -8.264 | -6.418 | -4.607 | -5.055 | -4.073 | -2.422 |
| marR | 24 | -8.438 | -5.887 | -4.483 | -5.111 | -4.072 | -2.932 |
| matA | 56 | -8.966 | -3.892 | -3.201 | -3.654 | -2.609 | -1.619 |
| mdetl1 | 8 | -4.835 | -4.087 | -3.899 | -3.979 | -3.726 | -3.496 |
| mdtA | 9 | -9.770 | -6.711 | -4.845 | -5.651 | -3.883 | -3.146 |
| mdtEyhiU | 59 | -7.460 | -4.561 | -3.779 | -4.145 | -3.304 | -2.464 |
| mecA | 31 | -7.921 | -6.533 | -4.523 | -4.738 | -3.373 | -0.222 |
| mefA | 121 | -9.925 | -3.617 | -2.691 | -2.897 | -1.519 | -0.255 |

| | | | | | | | |
|---|---|---|---|---|---|---|---|
| mepA | 64 | -7.887 | -3.839 | -3.523 | -3.823 | -3.213 | -1.425 |
| merA | 2 | -7.273 | -7.249 | -7.224 | -7.224 | -7.200 | -7.176 |
| mexA | 17 | -8.726 | -7.456 | -5.182 | -5.470 | -3.799 | -1.294 |
| mexB | 14 | -6.693 | -3.993 | -3.915 | -4.209 | -3.698 | -3.440 |
| mexE | 42 | -8.263 | -5.474 | -4.385 | -4.921 | -3.914 | -2.845 |
| mexF | 89 | -6.144 | -2.635 | -2.225 | -2.469 | -1.785 | -1.233 |
| mcr1 | 154 | -8.495 | -6.816 | -5.689 | -5.605 | -4.730 | -2.063 |
| mphA | 116 | -6.966 | -4.617 | -2.908 | -3.460 | -2.429 | -1.619 |
| mphB | 28 | -8.919 | -4.960 | -4.440 | -4.909 | -3.774 | -2.520 |
| mphC | 12 | -9.215 | -4.899 | -4.495 | -5.126 | -4.206 | -3.597 |
| msrE | 25 | -4.786 | -3.409 | -2.895 | -2.751 | -2.241 | -0.100 |
| msrA | 24 | -6.448 | -4.605 | -4.347 | -4.373 | -3.818 | -3.340 |
| msrC | 31 | -8.775 | -6.340 | -4.313 | -4.933 | -3.829 | -1.181 |
| mtrD | 56 | -9.798 | -5.424 | -3.804 | -4.522 | -3.402 | -2.285 |
| nimE | 8 | -4.869 | -4.469 | -4.310 | -4.239 | -4.057 | -3.577 |
| nisB | 13 | -5.978 | -5.228 | -4.531 | -4.562 | -3.800 | -3.534 |
| oleC | 42 | -7.673 | -4.245 | -3.330 | -4.012 | -3.140 | -2.532 |
| oprD | 86 | -8.327 | -5.273 | -3.283 | -3.914 | -2.623 | -1.130 |
| oprJ | 97 | -9.798 | -3.670 | -2.947 | -3.329 | -2.448 | -1.613 |
| oqxA | 12 | -8.493 | -8.129 | -7.694 | -7.667 | -7.481 | -6.385 |
| oqxB | 15 | -5.787 | -4.478 | -3.972 | -3.601 | -2.960 | 0.197 |
| pbp | 10 | -5.312 | -4.097 | -2.430 | -3.109 | -2.327 | -2.130 |
| pbp5 | 12 | -4.809 | -4.251 | -3.937 | -4.011 | -3.696 | -3.476 |
| pcoA | 1 | -0.285 | -0.285 | -0.285 | -0.285 | -0.285 | -0.285 |
| Pe | 10 | -4.184 | -3.588 | -3.403 | -3.422 | -3.058 | -2.936 |
| penA | 38 | -7.799 | -6.408 | -4.440 | -4.820 | -3.615 | -2.775 |
| pica | 32 | -6.140 | -5.255 | -4.792 | -4.305 | -3.074 | -2.373 |
| pikR1 | 13 | -9.353 | -7.887 | -7.733 | -6.357 | -4.000 | -3.252 |
| pikR2 | 49 | -7.662 | -4.172 | -3.552 | -4.029 | -3.252 | -2.842 |
| pmrA | 9 | -5.473 | -3.831 | -3.763 | -3.927 | -3.585 | -3.137 |
| pncA | 49 | -7.757 | -3.814 | -3.257 | -3.633 | -2.851 | -2.178 |
| qacA | 7 | -8.680 | -6.829 | -4.404 | -5.586 | -4.331 | -3.698 |
| qacF | 2 | -6.537 | -6.395 | -6.253 | -6.253 | -6.112 | -5.970 |
| qacH | 70 | -8.232 | -4.570 | -3.262 | -3.842 | -2.584 | -1.645 |
| qepA | 27 | -3.615 | -2.793 | -2.436 | -2.156 | -1.578 | 0.197 |
| qnrA | 136 | -8.369 | -6.554 | -4.911 | -4.935 | -3.414 | -1.176 |
| qnrB | 26 | -5.653 | -4.350 | -3.424 | -3.586 | -2.824 | -1.971 |
| qnrD | 41 | -7.673 | -6.424 | -4.784 | -4.767 | -2.996 | -1.888 |
| qnrS | 386 | -9.100 | -4.993 | -4.058 | -4.117 | -3.032 | 0.471 |
| rarD | 22 | -8.752 | -4.034 | -3.601 | -4.134 | -3.195 | -2.573 |
| rcnA | 2 | -7.489 | -7.456 | -7.423 | -7.423 | -7.390 | -7.357 |
| sat4 | 27 | -4.719 | -4.080 | -3.564 | -3.450 | -2.826 | -1.918 |
| sdeB | 17 | -7.813 | -3.939 | -3.727 | -4.183 | -3.433 | -3.142 |

| | | | | | | | |
|---|---|---|---|---|---|---|---|
| spcN | 26 | -8.967 | -6.927 | -5.082 | -5.464 | -4.092 | -3.124 |
| str | 18 | -4.427 | -3.906 | -3.319 | -3.254 | -2.687 | -2.000 |
| strA | 52 | -9.145 | -4.483 | -3.787 | -3.492 | -1.738 | -0.363 |
| strB | 111 | -8.619 | -3.598 | -2.667 | -2.965 | -1.891 | -0.565 |
| sul1 | 786 | -8.926 | -3.185 | -2.181 | -2.258 | -1.227 | 4.800 |
| sul2 | 590 | -9.115 | -3.515 | -2.593 | -2.650 | -1.592 | 1.826 |
| sul3 | 51 | -5.640 | -5.124 | -3.036 | -3.303 | -2.140 | 0.456 |
| sulA | 39 | -8.840 | -6.295 | -5.270 | -5.607 | -4.718 | -3.233 |
| tet32 | 53 | -4.426 | -3.212 | -2.799 | -2.858 | -2.536 | -1.863 |
| tet34 | 57 | -9.165 | -4.670 | -4.136 | -4.397 | -3.678 | -1.266 |
| tet35 | 1 | -4.588 | -4.588 | -4.588 | -4.588 | -4.588 | -4.588 |
| tet36 | 19 | -4.847 | -4.055 | -3.887 | -3.829 | -3.608 | -2.130 |
| tet37 | 13 | -8.762 | -7.955 | -4.458 | -5.114 | -2.727 | -1.328 |
| tet38 | 1 | -4.426 | -4.426 | -4.426 | -4.426 | -4.426 | -4.426 |
| tetA | 399 | -8.958 | -3.481 | -1.811 | -2.125 | -0.414 | 1.858 |
| tetB | 159 | -8.796 | -4.022 | -3.362 | -3.352 | -2.394 | 0.485 |
| tetD | 53 | -8.815 | -6.517 | -4.626 | -5.083 | -3.817 | -1.844 |
| tetE | 70 | -8.780 | -5.658 | -4.021 | -4.398 | -2.983 | -1.161 |
| tetG | 364 | -8.394 | -3.157 | -2.297 | -2.256 | -1.125 | 2.201 |
| tetH | 183 | -9.828 | -3.223 | -2.025 | -2.226 | -0.759 | 1.040 |
| tetJ | 4 | -5.004 | -4.235 | -3.959 | -4.199 | -3.923 | -3.874 |
| tetk | 19 | -8.535 | -3.202 | -2.723 | -3.238 | -2.603 | -1.738 |
| tetL | 89 | -9.590 | -3.958 | -3.073 | -3.379 | -2.372 | -0.153 |
| tetM | 420 | -9.119 | -4.417 | -2.968 | -3.285 | -2.117 | 1.092 |
| tetO | 346 | -10.000 | -3.442 | -2.159 | -1.849 | 0.347 | 2.150 |
| tetPB | 61 | -8.726 | -4.744 | -3.862 | -4.254 | -3.366 | -2.672 |
| tetQ | 326 | -8.071 | -2.765 | -1.443 | -1.491 | 0.581 | 2.235 |
| tetS | 48 | -4.964 | -4.044 | -3.333 | -3.329 | -2.870 | -1.091 |
| tetT | 26 | -5.116 | -3.971 | -3.647 | -3.362 | -2.896 | -0.840 |
| tetU | 4 | -8.633 | -7.777 | -7.481 | -7.150 | -6.854 | -5.007 |
| tetV | 5 | -4.835 | -4.567 | -4.280 | -4.210 | -4.140 | -3.228 |
| tetW | 352 | -8.175 | -3.903 | -2.803 | -3.012 | -2.072 | 0.678 |
| tetX | 273 | -9.661 | -3.745 | -2.515 | -2.811 | -1.831 | 0.701 |
| tetZ | 26 | -5.926 | -5.404 | -3.416 | -3.820 | -2.544 | -0.242 |
| tet39 | 7 | -3.990 | -3.015 | -2.190 | -2.097 | -1.375 | 0.280 |
| tetC | 108 | -5.480 | -3.599 | -2.772 | -2.549 | -1.677 | 0.640 |
| tetPA | 27 | -4.300 | -3.870 | -3.570 | -3.515 | -3.246 | -2.584 |
| tetR | 89 | -7.469 | -4.493 | -3.279 | -3.774 | -2.936 | -1.676 |
| tnpA1 | 95 | -6.606 | -4.432 | -3.232 | -3.042 | -2.258 | -0.114 |
| tnpA2 | 65 | -5.969 | -3.697 | -3.293 | -3.197 | -2.487 | -1.114 |
| tnpA3 | 64 | -5.932 | -4.123 | -3.433 | -3.266 | -2.338 | -0.382 |
| tnpA4 | 86 | -5.506 | -3.727 | -2.704 | -2.781 | -1.791 | -0.772 |
| tnpA5 | 69 | -5.398 | -3.593 | -2.629 | -2.711 | -1.760 | -0.770 |

| | | | | | | | |
|---|---|---|---|---|---|---|---|
| tnpA6 | 38 | -5.622 | -4.100 | -3.101 | -3.191 | -2.657 | -1.169 |
| tnpA7 | 59 | -5.400 | -3.224 | -2.614 | -2.754 | -2.163 | -1.178 |
| tolC | 55 | -8.140 | -4.193 | -3.752 | -4.104 | -3.371 | -2.597 |
| Tp614 | 44 | -6.185 | -4.347 | -3.323 | -3.400 | -2.334 | -0.870 |
| ttgA | 32 | -8.733 | -4.893 | -4.181 | -4.872 | -3.930 | -3.137 |
| ttgB | 41 | -8.650 | -5.741 | -4.560 | -5.012 | -3.950 | -2.976 |
| uidA | 5 | -4.915 | -4.574 | -3.594 | -3.649 | -3.573 | -1.590 |
| vanA | 26 | -4.969 | -3.297 | -2.390 | -2.316 | -1.255 | 0.262 |
| vanB | 65 | -7.869 | -4.000 | -3.550 | -3.991 | -3.188 | -2.155 |
| vanC | 70 | -7.824 | -3.665 | -2.849 | -3.391 | -2.660 | -2.167 |
| vanG | 3 | -4.488 | -4.338 | -4.188 | -4.218 | -4.083 | -3.979 |
| vanHB | 42 | -7.754 | -4.176 | -3.689 | -4.134 | -3.354 | -2.195 |
| vanHD | 5 | -8.422 | -8.256 | -7.915 | -6.630 | -4.313 | -4.243 |
| vanRA | 9 | -4.626 | -4.347 | -4.149 | -4.063 | -3.955 | -2.567 |
| vanRB | 9 | -8.314 | -7.740 | -5.411 | -6.059 | -4.458 | -4.117 |
| vanRC | 2 | -5.587 | -5.510 | -5.434 | -5.434 | -5.358 | -5.281 |
| vanSB | 26 | -8.592 | -4.855 | -4.209 | -4.879 | -3.972 | -2.349 |
| vanSC | 12 | -9.542 | -8.302 | -4.776 | -6.000 | -4.115 | -3.653 |
| vanTC | 25 | -8.416 | -7.342 | -4.180 | -5.244 | -3.862 | -3.453 |
| vanTE | 2 | -8.674 | -7.495 | -6.316 | -6.316 | -5.137 | -3.959 |
| vanTG | 4 | -8.644 | -5.707 | -4.631 | -5.555 | -4.478 | -4.313 |
| vanWB | 4 | -9.668 | -5.842 | -4.253 | -5.390 | -3.801 | -3.387 |
| vanWG | 19 | -7.110 | -4.318 | -3.748 | -3.639 | -2.800 | -1.400 |
| vanXA | 2 | -9.228 | -9.057 | -8.886 | -8.886 | -8.716 | -8.545 |
| vanXD | 14 | -5.665 | -5.312 | -5.029 | -4.639 | -3.944 | -2.716 |
| vanYB | 10 | -9.841 | -9.079 | -7.262 | -6.992 | -4.725 | -4.101 |
| vanYD | 31 | -8.438 | -5.678 | -4.117 | -4.845 | -3.677 | -3.213 |
| vatB | 2 | -4.456 | -3.914 | -3.373 | -3.373 | -2.831 | -2.290 |
| vatC | 6 | -4.626 | -4.549 | -4.390 | -4.414 | -4.276 | -4.241 |
| vatD | 2 | -3.831 | -3.669 | -3.508 | -3.508 | -3.346 | -3.184 |
| vatE | 50 | -9.726 | -5.676 | -4.413 | -4.855 | -3.773 | -1.313 |
| vgaA | 14 | -9.969 | -7.657 | -4.702 | -5.894 | -4.166 | -3.395 |
| vgaB | 24 | -7.932 | -6.673 | -4.374 | -4.970 | -3.901 | -2.636 |
| yceEmdtG1 | 14 | -4.969 | -3.827 | -3.718 | -3.574 | -2.984 | -2.547 |
| yceLmdtH1 | 38 | -8.445 | -4.177 | -3.719 | -4.305 | -3.414 | -2.652 |
| yceLmdtH2 | 18 | -5.894 | -4.602 | -4.130 | -4.070 | -3.681 | -2.650 |

**Table S2. Results of the LMM models (‘year’ was set as a fixed factor and ‘country’ as a random factor, the test performed on max values).**

| Environment | Effect size | p-value | Bonferroni |
|---|---|---|---|
| biofilm | 0.4142 | 0.0000055 | 0.000055 |
| effluent | -0.10799 | 0.000994 | 0.00994 |
| feces | 0.3117 | 0.195 | 1 |
| food | 1.4472 | 2E-16 | 2E-15 |
| manure | 0.02423 | 0.1163 | 1 |
| sediments | -0.10465 | 6.59E-09 | 6.59E-08 |
| sludge | -0.305 | 4.61E-11 | 4.61E-10 |
| soil | 0.02474 | 0.0241 | 0.241 |
| wastewater | -0.25085 | 1.69E-12 | 1.69E-11 |
| water | -0.15385 | 0.0106 | 0.106 |

**Figure S3. Correlation analysis between the selected ARGs.**

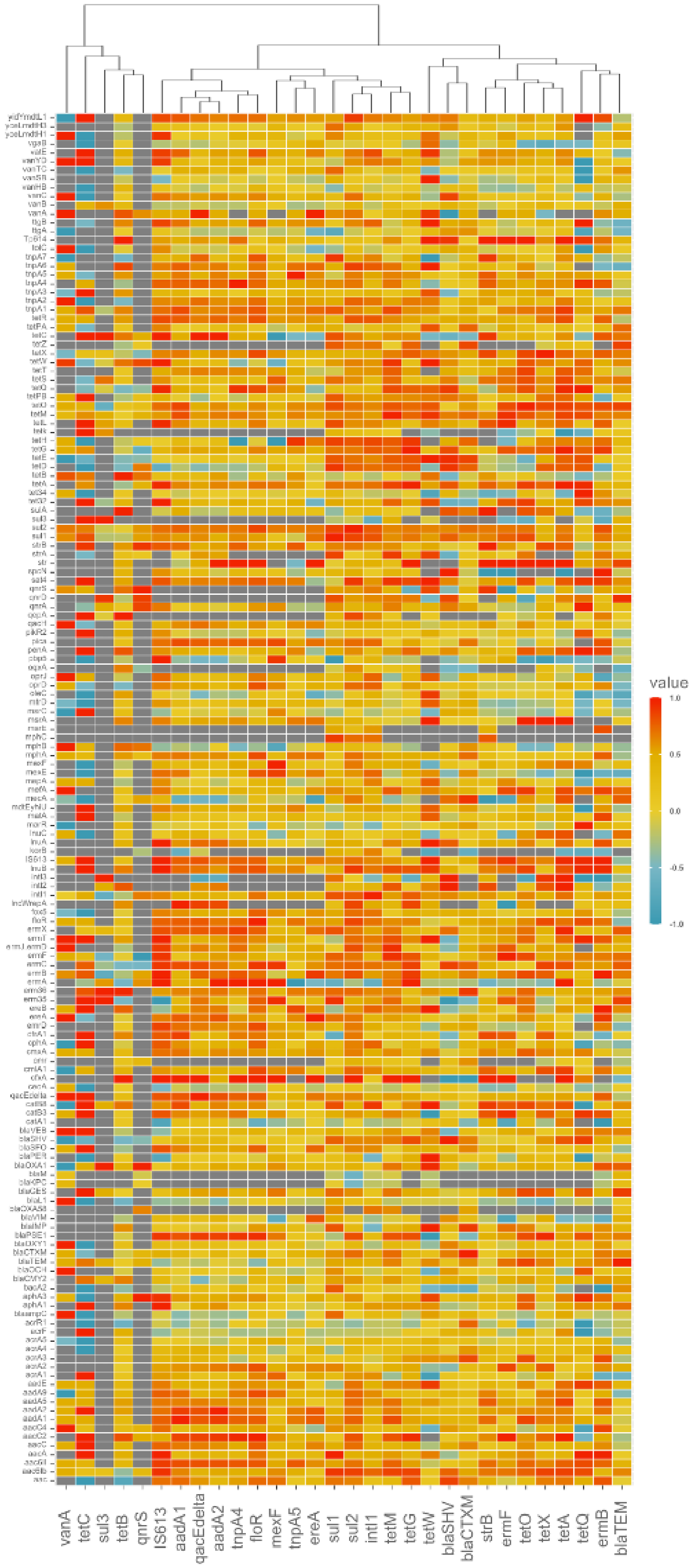

**Figure S4. Relationship between ARG abundances and time: the data points are the abundances of ARGs in samples from each country and the lines are fitted to the maximum abundance values; * denotes significant trends.**

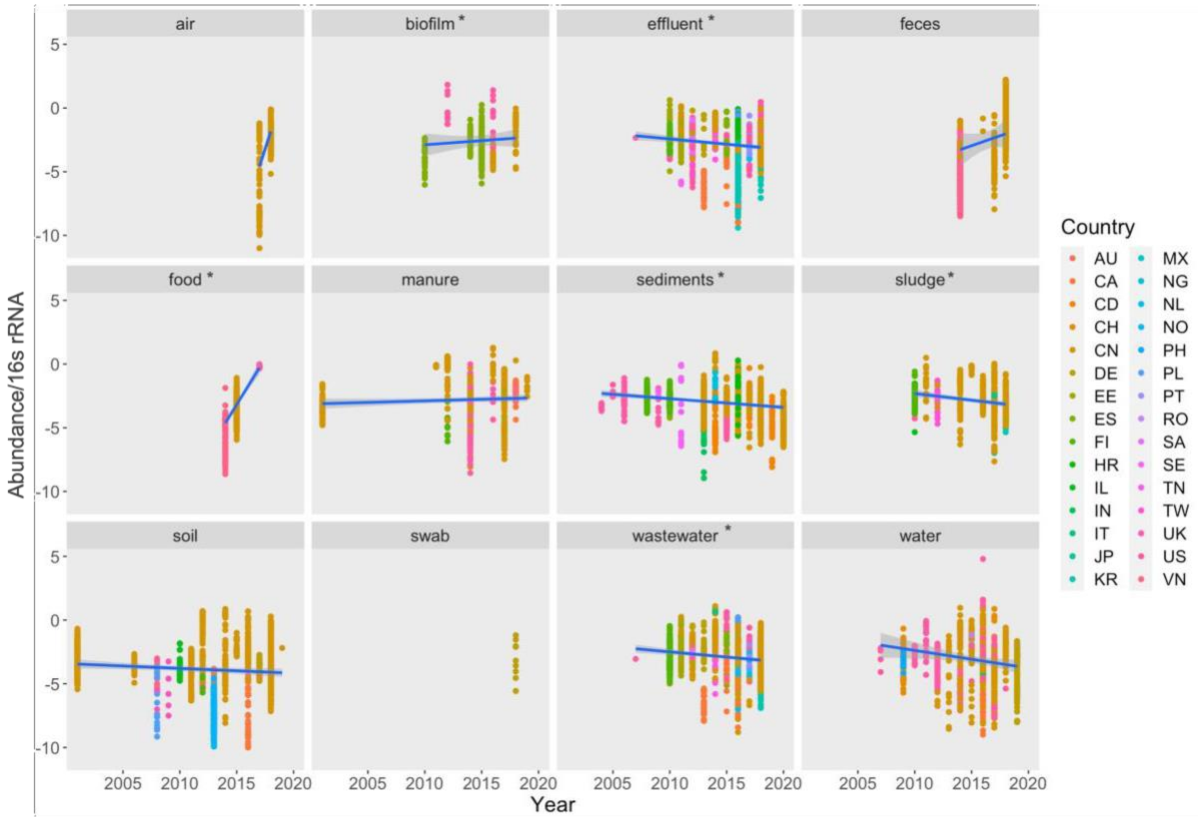

**Figure S5. Abundance of most reported ARGs. ARGs reported in more than 20 studies are represented in orange, and ARGs reported less but which were generally highly abundant are shown in blue. The line represents an average abundance across all data points.**

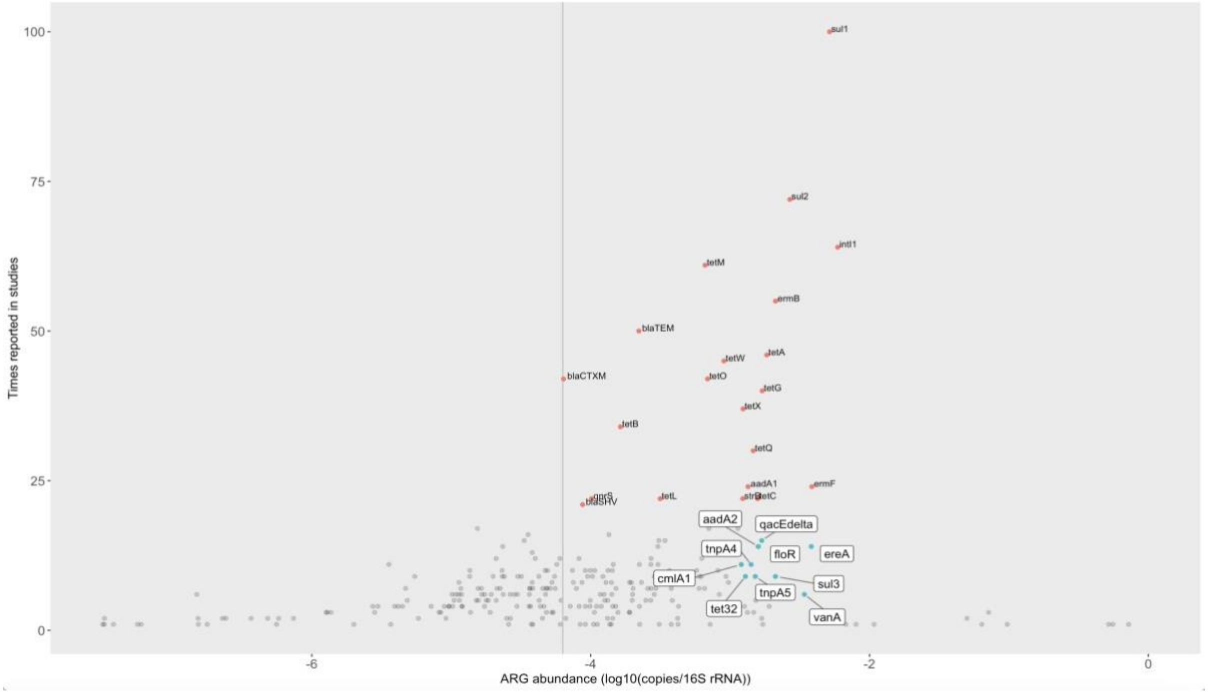

**Figure S6. Density plots represent abundance ranges of the most reported ARGs in different environmental types divided according to exposure to anthropogenic impact (see material and methods for the detailed description), and corresponding bar plots show what samples contributed to each distribution. The environmental types are color-coded as follows: "feces/manure" in purple, "impacted" in orange, "likely impacted" in yellow, "likely unimpacted" in green and "unknown" type in blue. Note that abundances are given relative to the number of 16S rRNA copies, which means that these distributions do not necessarily correspond to the total exposure to resistant bacteria in a given environment.**

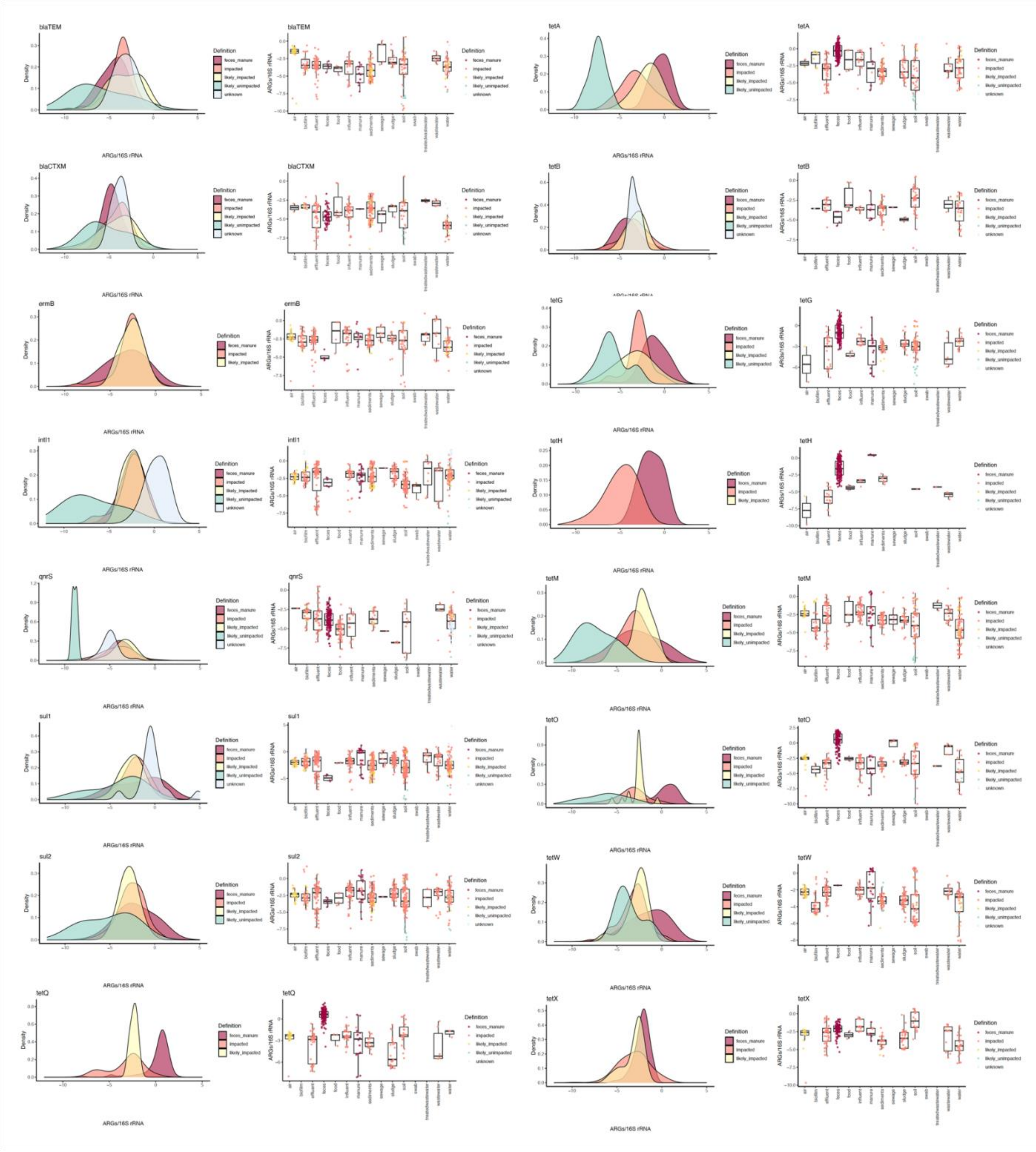
